# Complement Factor H and its *C. elegans* homolog regulate IFT52/OSM-6 and CNG channel localization in sensory neurons

**DOI:** 10.1101/2025.10.10.681644

**Authors:** Yumiko Oshima, Katarzyna A. Hussey, Joanna Hagen, Zeljka Smit-McBride, Maya Moorthy, Joanne A. Matsubara, Martin Flajnik, Robert J. Johnston, Bruce E. Vogel

## Abstract

Age-related macular degeneration (AMD), the leading cause of blindness in the elderly, is characterized by progressive degeneration of retinal photoreceptors. Current disease models propose AMD pathogenesis is a consequence of cytolytic damage and tissue inflammation that result from defective repression of alternative complement pathway activity by complement factor H (CFH). However, recent studies demonstrate functions for CFH that are outside of its established role in the alternative complement pathway, suggesting that novel CFH-mediated mechanisms may influence AMD initiation and progression. Our previous demonstration that CFH and its nematode homolog, CFH-1, modulate inversin/NPHP-2 accumulation in vertebrate photoreceptor and *C. elegans* sensory neuron cilia during aging suggests that AMD patients with CFH loss-of-function mutations have cilia defects that may contribute to photoreceptor dysfunction. Here, we investigate the consequences of CFH and CFH-1 loss-of-function mutations on the dynamics and localization of intraflagellar transport (IFT) train and visual cycle components in these cells. In *C. elegans* sensory neurons, IFTB1 components IFT52/OSM-6 and IFT88/OSM-5 are transported at similar rates in WT animals but IFT52/OSM-6 transport slows significantly in *cfh-1* mutant animals while IFT88/OSM-5 is unaffected. Defective localization of IFT52/OSM-6 in photoreceptors of CFH knockout mice and in human photoreceptors from AMD high-risk CFH Y402H homozygotes, suggest an evolutionarily conserved role for CFH in promoting IFT52/OSM-6 transport and localization in sensory neuron cilia. In addition, distribution of CNG channel subunits in *C. elegans cfh-1* mutant sensory neurons and CFH Y402H high-risk human photoreceptors are distinct from their WT and Y402 low-risk counterparts. Together, the data indicate previously unappreciated functions for CFH in IFT train organization and cilia protein localization and suggest a novel mechanism for photoreceptor segment thinning, an early AMD biomarker that has been linked to CFH high-risk variants.

## Introduction

Age-related macular degeneration (AMD) is a leading cause of blindness among the elderly, affecting 11% of adults over the age of 85, although the mechanisms of disease initiation and progression are not fully understood. Defective regulation of the alternative complement pathway, part of the innate immune system’s natural defense against infections, was implicated as a key contributor to AMD progression by the identification of amino acid substitutions (Y402H and R1210C) in complement factor H (CFH) as AMD risk factors (1-6). CFH is a secreted protein that negatively regulates C3 activity by promoting decay of the C3bBb convertase that activates C3 and by acting as a cofactor for factor I protease in processing C3b to its inactive form (7). Current models for AMD suggest that the accumulation of drusen, extracellular deposits located between the retinal pigment epithelium (RPE) and Bruch’s membrane, coupled with defective regulation of C3 by CFH, result in inflammation and cytolysis that disrupt the function of RPE cells and their glial-like interactions with photoreceptors (8,9).

Early AMD biomarkers that appear well in advance of established clinical symptoms may give insight into the initial stages of AMD pathogenesis (10-12). Among these biomarkers, photoreceptor segment (PS) thinning and defects in rod-mediated dark adaptation (RMDA) have been linked to high-risk CFH variants although the mechanistic links between established CFH functions and these pre-clinical biomarkers are not obvious (10,11). Defective RMDA is detected in regions within the retina that coincide with defects in interdigitation zone (IZ) integrity, indicating the possibility that these two biomarkers may share a common mechanism (12). Recent studies have shown that CFH has novel functions in monocyte migration, lipid distribution, and cilia compartment organization, suggesting that mechanisms outside the alternative complement pathway may contribute to the appearance of early AMD biomarkers and the mechanisms of disease initiation and progression (13-15).

Vertebrate photoreceptors are highly polarized cells with structurally distinct regions that include inner segments where the cellular components of phototransduction are synthesized and outer segments where the phototransduction cascade is initiated (16). Between these regions is a connecting cilium composed of a microtubule based axoneme that extends into the outer segments and contains the machinery for the intraflagellar transport (IFT) of phototransduction components between inner and outer segments (17, 18). Although *C. elegans* do not have a visual system with opsin-based photoreceptors, 60 ciliated sensory neurons detect chemical, thermal, and mechanical stimuli (19, 20). Like their vertebrate counterparts, nematode cilia are microtubule-based cell protrusions with IFT machinery and structurally distinct cilia compartments that are highly conserved between species (20-22). In between the proximal transition zone (TZ) and the cilia distal segment (DS) is the middle segment (MS) with a domain called the “inversin compartment” defined by the location of the inversin protein (also called NPHP-2) in many cilia subtypes (23, 24). In a previous study, we demonstrated that loss-of-function mutations in the *C. elegans* CFH homolog, CFH-1, results in progressive inversin accumulation and defective cilia function in the sensory neurons of aging animals (15). Similar defects in inversin/NPHP-2 distribution were also detected in CFH knockout mice and human photoreceptors from individuals homozygous for the AMD high-risk CFH Y402H variant, suggesting that cilia structural defects make a significant, yet unappreciated contribution to the progressive photoreceptor dysfunction observed in AMD patients with CFH loss-of-function mutations (15).

Here, we examine the consequences of CFH-1 and CFH loss of function on the distribution of evolutionarily conserved proteins involved in intraflagellar transport (IFT) and the vertebrate visual cycle and find that CFH and its invertebrate homolog are required for IFT52/OSM-6 and cyclic nucleotide gated (CNG) channel localization or transport in vertebrate photoreceptors and *C. elegans* ciliated chemosensory neurons. Together, our data support a model that CFH loss of function variants contribute to AMD due, in part, to defects in photoreceptor cilia organization and protein transport and suggest IFT-based defects may contribute to early AMD biomarkers associated with high-risk CFH variants.

## RESULTS

### *C. elegans cfh-1* mutant animals have defects in intraflagellar transport (IFT)

Intraflagellar transport (IFT) trains are large evolutionarily conserved polymers of IFT-A and IFT-B protein complexes that transport structural and signaling cargo from cilia base to tip (i.e. anterograde transport) and from tip to base (i.e. retrograde transport) using kinesin and dynein motors (25). Based on the distribution of CFH-1 in close proximity to sensory neuron cilia and the disruption of inversin/NPHP-2 distribution in aging adult *cfh-1* mutants described previously (15), in addition to the observation that another component of the NPHP module, NPHP-4, negatively regulates IFT rates (26), we compared the movement of several IFT train components in the phasmid neurons of WT and *cfh-1* mutant *C. elegans* day 4 adults. Most of the components examined had small, but statistically significant, increases in the rate of anterograde transport in *cfh-1(em14)* adult mutant animals when compared to age-matched WT animals (Figs. 1, S1). These included dynein light-intermediate chain XBX-1, IFT-A component IFT140/CHE-11, and IFTB1 component IFT88/OSM-5. [*n*.*b*. IFT rates shown in Figures 1 and S1 for WT day 4 adult animals are slower than published IFT rates for younger animals. This is likely because IFT has been shown to slow with age (27)]. In contrast to IFT88/OSM-5, another IFTB1 component, IFT52/OSM-6, had a significantly slower rate of anterograde and retrograde transport in *cfh-1(em14)* day 4 adult mutant animals. The data suggest distinct effects of CFH-1 on IFT52/OSM-6 and IFT88/OSM-5 rates even though both are subunits of the IFTB1 complex (25).

**Figure 1.**
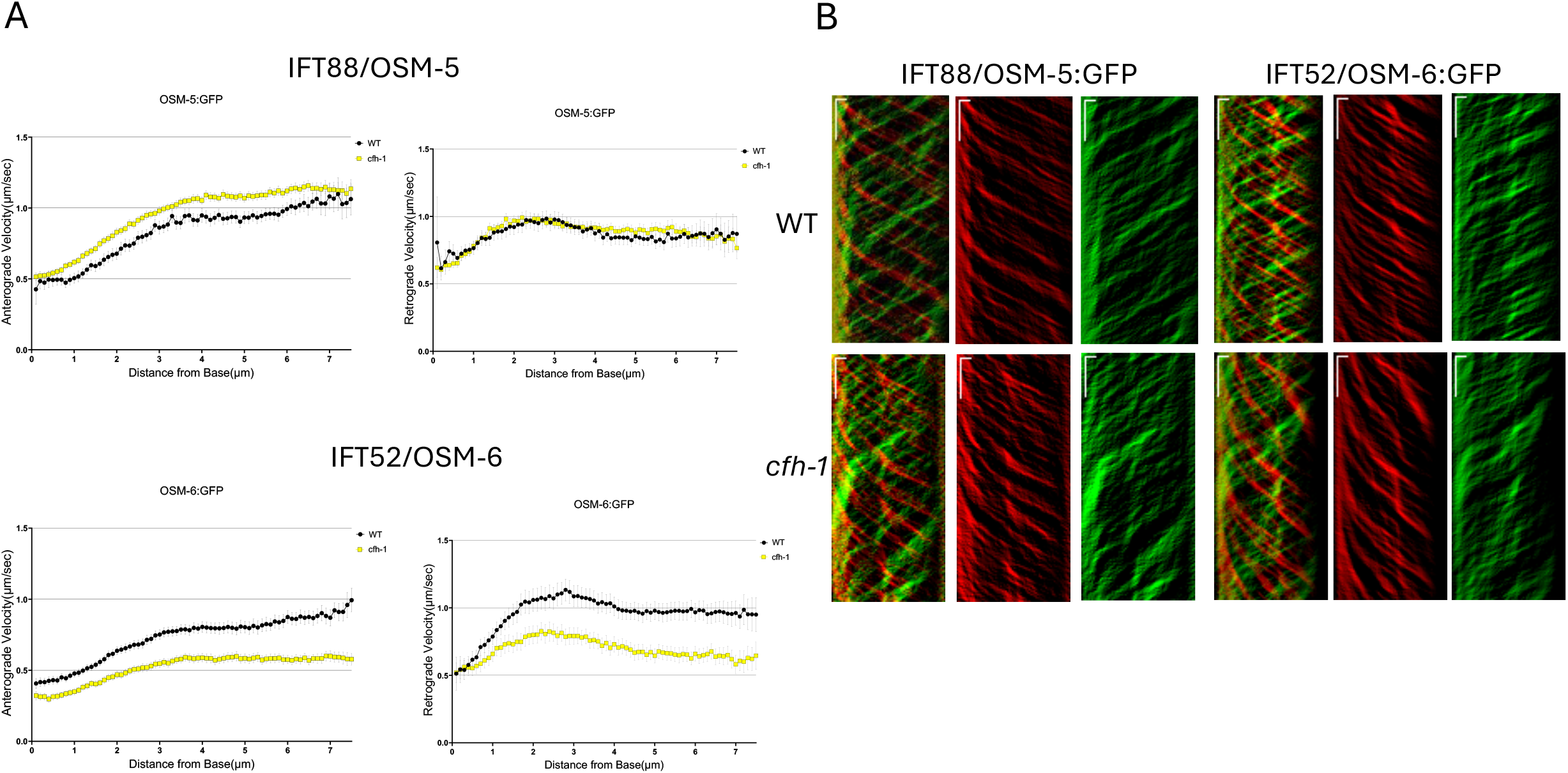
Comparison of anterograde and retrograde movement of IFT88/OSM-5 and IFT52/OSM-6 in WT and cfh-1 mutant C. elegans day 4 adults. A, Plots of anterograde (left) and retrograde (right) IFT velocity vs. distance from cilia base for IFT88/OSM-5::GFP (top) and IFT52/OSM-6::GFP (bottom) in phasmid sensory neurons of WT(black) and *cfh-1* mutant (yellow) animals. Note that in *cfh-1* mutant animals IFT velocities for IFT88/OSM-5 are similar to WT in retrograde IFT and slightly faster in anterograde IFT. In contrast, IFT velocities are decreased significantly for another IFTB component, IFT52/OSM-6, in both directions in *cfh-1* mutant animals. B, Representative anterograde (red) and retrograde (green) kymograph plots upon which the data are based. Horizonal scale bars represent 1µm and vertical scale bars indicate a 5 second interval.

### *C. elegans* CFH-1 effects on IFT are not a result of inversin/NPHP-2 gain-of-function

In previous work, we demonstrated that inversin/NPHP-2 localizes within its namesake proximal cilia compartment in WT animals, but accumulates in distal cilia compartments in CEP sensory neurons of aging *cfh-1* mutant animals (15). Similar ectopic accumulation of inversin/NPHP-2 was observed in the photoreceptors of cfh knockout mice and of human homozygous for the high AMD-risk Y402H variant (15). To determine whether the abnormal rate of IFT-52/OSM-6 IFT in *cfh-1* day 4 adult mutant phasmid neurons is due to ectopic increase in inversin/NPHP-2 function, IFT52/OSM-6 IFT rates were first compared in age-matched WT and *nphp-2(gk653)* single mutants (Fig. S2A). Next, IFT rates were compared in *cfh-1(em14)* single mutants and *cfh-1(em14); nphp-2(gk653)* double mutants (Fig. S2B). In both cases IFT52/OSM-6 IFT rates were not affected by *nphp-2* loss-of-function, suggesting that increased NPHP-2 activity is not responsible for the decreased rate of IFT52/OSM-6 IFT in *cfh-1* mutant animals(Fig. S2).

### CFH localizes preferentially on human rod photoreceptor outer segments

Previous demonstrations of CFH localization on mouse photoreceptors (15,28,29) coupled with inversin compartment defects in AMD high-risk CFH Y402H photoreceptors (15) and the link of CFH variants with RMDA defects and photoreceptor segment thinning (10) raised questions regarding CFH distribution on human photoreceptors. To determine CFH localization on human photoreceptors, retinal organoids generated from human stem cells were examined with CFH-specific antibodies in tandem with rod specific rhodopsin and cone-specific opsin antibodies (Fig. S3). A significantly higher proportion of rhodopsin-positive photoreceptors were also positive for CFH [51.6% (116/225)] compared to cones where only 1.8% (4/225) were positive for CFH (Fig. S3)]. Interestingly, in these organoids CFH protein distribution appears to be primarily internal to rhodopsin staining at the photoreceptor periphery (Fig. S3).

To identify CFH localization on aged adult photoreceptors, postmortem retina sections from individuals over 50 years old were stained with CFH antibodies. In contrast to previously observed CFH staining of mouse retina on photoreceptor inner segments (15,28), CFH staining of human retina had extensive overlap with rhodopsin on photoreceptor outer segments (Fig. S3).

### IFT52/OSM-6 distribution is defective in CFH knockout mouse and AMD high-risk CFH Y402H human photoreceptors

To determine whether the CFH-1 dependent IFT defects described above for *C. elegans* sensory neurons (Fig. 1) apply to IFT components in vertebrate photoreceptors, we compared IFT52/OSM-6, IFT88/OSM-5, and rhodopsin localization in retinas from WT and cfh−/− knockout mice (21- and 19-wk-old males, respectively). IFT52/OSM-6 is found primarily in photoreceptor inner segments of WT and cfh-/-mice. However, there is a statistically significant difference in the ratio of IFT52/OSM-6 in photoreceptor inner and outer segments between WT and CFH knockout mice (Fig. 2). There was also a small difference in IFT88/OSM-5 staining between WT and cfh -/-knockout mice that was not statistically significant and no detectable difference in rhodopsin localization. This suggests that in parallel to the role of CFH-1 in transport of IFT52/OSM-6 in *C. elegans* sensory neurons, CFH is necessary for the correct distribution of IFT52/OSM-6 in mouse photoreceptors.

**Figure 2.**
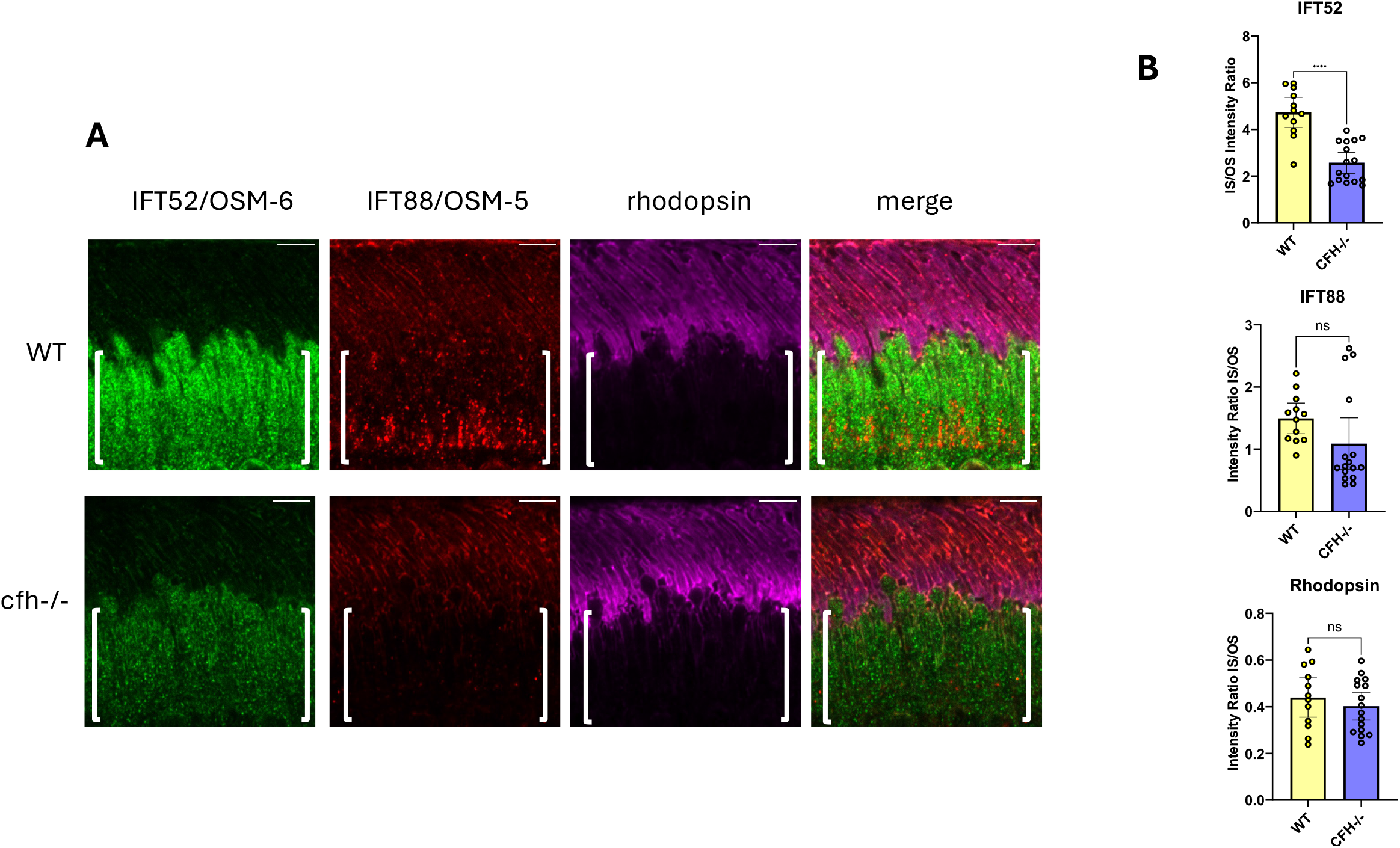
Mislocalization of IFT-52/OSM-6 in cfh knockout mouse photoreceptors. A, *Comparison of IFT-52/OSM-6, IFT-88/OSM-5, and rhodopsin localization in WT and cfh knockout mouse photoreceptors*. Retina sections from WT and cfh knockout mice were stained with antibodies for IFT52/OSM-6, IFT88/OSM-5, and rhodopsin. Note the decreased amount of IFT52/OSM-6 staining in inner segments (indicated by brackets) of cfh knockout mice in comparison to WT mice. Scale bars are 5µm. Posterior is oriented up in all images. B, scatter plots of relative fluorescence levels in inner segment divided by outer segment for IFT52/OSM-6, IFT88/OSM-5, and rhodopsin in WT and cfh knockout animals. Error bars indicate 95% confidence intervals. Significant difference from WT is indicated by brackets with asterisks as follows: **** P<.0001; ns indicates no significant difference from WT, P>.05.

The role of CFH in IFT52/OSM-6 distribution in mouse photoreceptors in combination with CFH distribution on outer segments of human photoreceptors (Fig. S3), raised the question of whether CFH affects IFT52/OSM-6 distribution in human photoreceptors. Postmortem retina sections from multiple AMD low-risk Y402 and high-risk Y402H individuals with no sign of disease [unrelated males between 52 and 68 y old (30)] were compared with respect to IFT52/OSM-6, IFT88/OSM-5, and rhodopsin localization. As in the mouse cfh -/- knockout (Fig. 2), human high-risk CFH Y402H photoreceptors appeared to have a decrease in the relative amount of IFT52/OSM-6 in photoreceptor inner segments that is apparent when comparing the ratio of inner segment to outer segment staining with IFT52/OSM-6 antibodies (Fig. 3). As in the mouse sections, small differences were also observed in the distribution of IFT88/OSM-5 and rhodopsin, however, these were not statistically significant.

**Figure 3.**
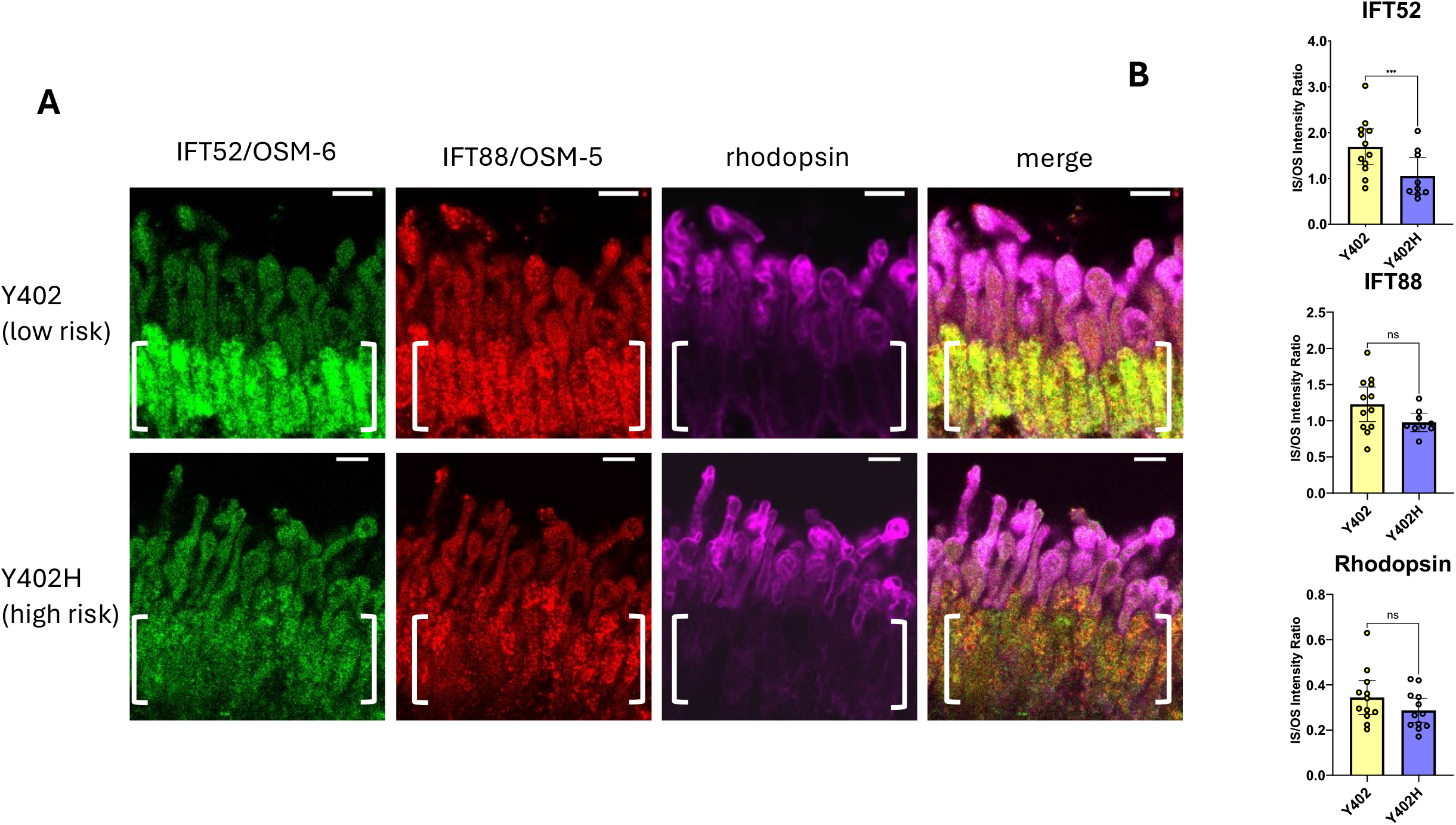
Mislocalization of IFT-52/OSM-6 in human Y402H high AMD risk retina. A, *Comparison of IFT-88/OSM-5, IFT-52/OSM-6, and rhodopsin localization in AMD Y402 low-risk and Y402H high-risk human photoreceptors*. Post-mortem retina sections from AMD Y402 low-risk and Y402H high-risk individuals were stained with antibodies for IFT52/OSM-6, IFT88/OSM-5, and rhodopsin. Note the decreased amount of IFT52/OSM-6 staining in inner segments of *AMD Y402H high-risk photoreceptors (brackets)*. Scale bars are 5µm. Posterior is oriented up in all images. B, scatter plots of relative fluorescence levels for IFT52/OSM-6, IFT88/OSM-5, and rhodopsin in inner segment divided by outer segment in photoreceptors from AMD Y402 low-risk and Y402H high-risk individuals. Error bars indicate 95% confidence intervals. Significant difference from WT is indicated by brackets with asterisks as follows: *** P<.001; ns indicates no significant difference from WT, P>.05.

### Disruption of CNG channel localization in *C. elegans cfh-1* mutant sensory neurons and human AMD high-risk photoreceptors

IFT is responsible for the transport of a number of components of the phototransduction cascade in photoreceptors (17). Since the distribution of IFT machinery is defective in CFH knockout mice and AMD high risk CFH Y402H human photoreceptors, it is important to know whether components of the phototransduction cascade are similarly affected. Previous studies have shown that the *C. elegans* cyclic nucleotide-gated channels CNGB/TAX-2 and CNGA/TAX-4 are located in the inversin compartment of amphid sensory neurons, and dependent on an intact compartment for correct distribution in a cell and subunit-specific manner (31,32). For example, in *nphp-2* mutant animals, CNGB/TAX-2 accumulates ectopically in the distal cilia compartment of AWB neurons, while CNGA/TAX-4 is unaffected. In contrast, CNGA/TAX-4 accumulates ectopically in the dendrite of ASK sensory neurons in *nphp-2* mutant animals while CNGB/TAX-2 is unaffected (31). Since mutations in *cfh-1* affect the integrity of the inversin compartment (15), CNGA/TAX-4 localization was compared in WT and *cfh-1* mutant backgrounds. In WT day 4 adults, TAX-4::GFP is localized in the dendrites and the inversin and distal compartments of phasmid cilia. In contrast, there was little or no detectable TAX-4::GFP in the distal compartment of *cfh-1* mutant phasmids, suggesting that in addition to inversin/NPHP-2 and OSM-6/IFT52, CFH-1 also regulates TAX-4 distribution in some *C. elegans* sensory neuron cilia (Fig. 4A).

**Figure 4.**
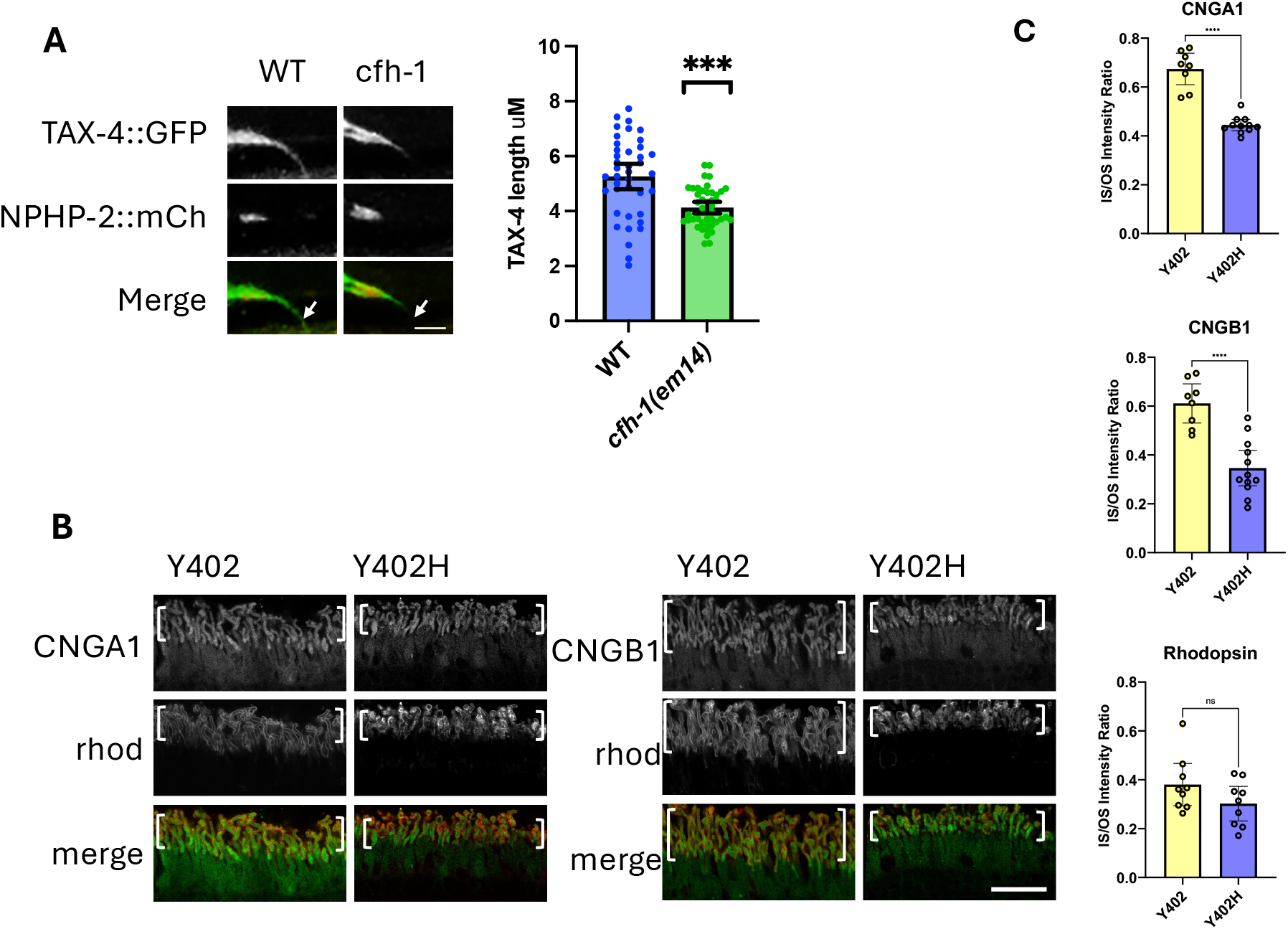
CFH-1 and CFH regulate CNG distribution in distal cilia compartments of C. elegans phasmid neurons and human photoreceptors. A, Comparison of TAX-4::GFP and inversin/NPHP-2::mCherry localization in phasmid neurons of WT *and cfh-1* mutant animals. Note relative lack of TAX-4::GFP in the distal cilia compartment of *cfh-1* mutant animals (arrows). Scale bars is 2.5µm. Right, scatter plots of TAX-4::GFP length in phasmid neurons of day 4 adult animals. Error bars indicate 95% confidence intervals. Significance is indicated by brackets with asterisks as follows: ***P<.001 B, Immunofluorescence with antibodies to CNGA1 (left) and CNGB1 in CFH Y402 AMD low-risk and CFHY402H AMD high-risk post-mortem human retina. Also shown is rhodopsin staining to indicate the location of photoreceptor outer segments (brackets). Scale bar is 20µm. C, scatter plots comparing relative fluorescence intensity in photoreceptor inner segments (IS) divided by outer segments (OS). Note the relative decrease in ratio of CNGA1 and CNGB1 fluorescence intensity in CFH Y402H AMD high risk photoreceptors. Significance is indicated by brackets with asterisks as follows: **** P<.0001; ns indicates no significant difference from WT, P>.05.

Based on this *cfh-1* mutant phenotype and the central role of CNG channels in the visual cycle in the vertebrate retina, the distribution of CNGA1 and CNGB1 were examined in post-mortem tissue from individuals at low-risk (CFH Y402) and high-risk (CFH Y402H) for AMD (30). A significant decrease in the relative quantity of CNGA1 and CNGB1 in inner segments when compared to outer segments was detected in AMD high-risk CFH Y402H variants (Fig. 4B,C), suggesting that CFH not only maintains inversin compartment boundaries (15), but also regulates the distribution of CNG channels in photoreceptor outer segments and that function is compromised in individuals at high-risk for AMD. Some CFH Y402H sections appear to show increased amounts of rhodopsin in photoreceptor outer segments, but measurements indicate that this increase is not statistically significant (Fig. 4B,C).

## DISCUSSION

The identification of CFH variants that significantly increase disease risk was a major advance in the AMD field and implicated defective inhibition of C3 and increased inflammation and cytolysis in disease pathogenesis. However, it is not obvious how ectopic C3 activity contributes to early AMD biomarkers linked to CFH high-risk variants that include photoreceptor layer thinning and delayed rod-mediated dark adaptation (RMDA) (10,11). Recent demonstrations of CFH functions that appear to be independent of its established role as a C3 inhibitor, suggest the existence of novel CFH mechanisms that may contribute to these early AMD biomarkers and the poorly understood early events of disease pathogenesis (13-15).

### A role for *cfh-1* in intraflagellar transport in *C. elegans* sensory neurons

Within the IFT train, IFT52/OSM-6 and IFT88/OSM-5 are components of the IFTB1 protein complex and behave similarly in WT animals, but very differently in aging *cfh-1* mutant animals in both anterograde and retrograde IFT trains (Fig. 1). This suggests that there may be some heterogeneity in IFTB1 composition in the absence of CFH-1. Interestingly, IFT52/OSM-6 assembly into retrograde IFT trains is unique in WT animals in that it is delayed compared to other IFT train components (33). The authors of that study suggest that IFT52/OSM-6 may be post-translationally modified before addition to retrograde trains (33). One possibility is CFH-1 facilitates this putative post-translational modification and promotes incorporation of IFT52/OSM-6 into the IFTB1 complex.

In previous work we identified a novel function for CFH and its nematode homolog in restricting inversin accumulation in the cilia of mouse and human photoreceptors and *C. elegans* sensory neurons (15). We tested whether inversin/NPHP-2 gain-of-function might affect IFT52/OSM-6 IFT by comparing its rate of transport in *cfh-1* single mutants and *cfh-1/nphp-2* double mutants but did not find a significant difference using an *nphp-2(gk653)* loss-of-function allele (34) (Fig. S2). This indicates that the observed slowing of IFT52/OSM-6 IFT in *cfh-1* mutant animals is not dependent on inversin/NPHP-2 gain-of-function, although it is possible that the *nphp-2(gk653)* allele produces truncated gene products that interfere with IFT52/OSM-6 function (34).

### CFH dependent IFT52/OSM-6 distribution in human and mouse photoreceptors and implications for photoreceptor layer thinning

Vertebrate photoreceptor cilia are highly dynamic, with approximately 800 molecules of rhodopsin transported from inner segments to outer segments every second (17). To determine whether there is a parallel to the role for *C. elegans* CFH-1 in IFT in vertebrate photoreceptors, IFT52/OSM-6 and IFT88/OSM-5 localization in mouse and human photoreceptors were analyzed in WT and CFH loss-of-function backgrounds (Figs. 2,3). In mouse and human tissue samples, loss of CFH function resulted in decreased relative amount of IFT52/OSM-6 within inner segments relative to outer segments (Figs. 2,3). This might reflect a decrease in retrograde trafficking and return of IFT52/OSM-6 to inner segments. As mentioned above for *C. elegans* CFH-1, CFH may promote the correct assembly of IFT52/OSM-6 into IFTB1 complexes in vertebrate retrograde IFT trains (17). Other possibilities are that the data reflect increased anterograde transport of IFT52/OSM-6 to photoreceptor outer segments or decreased IFT52/OSM-6 protein synthesis and/or trafficking to inner segments.

### Defective CNG channel distribution in human CFH Y402H photoreceptors

CFH is located on photoreceptor outer segments and preferentially associated with rods over cones in human tissue sections and retina organoids, respectively (Fig. 4), extending previous studies that demonstrated CFH localization in the photoreceptor layer of vertebrate retina (15, 28, 29).

The defect in localization of CNG channels previously seen in *C. elegans nphp-2* mutant cilia (31,32) suggests that inversin/NPHP-2 is necessary for the correct distribution of some molecular components essential to function of ciliated cells. Since disruption of inversin compartment integrity has been observed in *cfh-1* mutants (15), this raised the question of whether CNG channel distribution similarly depends on CFH-1 protein. The finding that TAX-4/CNGA distribution is significantly reduced in the distal segments of *cfh-1* mutant phasmid neurons suggests that the CFH-1 protein is involved in either the transport or maintenance of TAX-4/CNGA in the distal segments of sensory neuron cilia. It is possible that TAX-4/CNGA is directly dependent up IFT52/OSM-6, although that is difficult to test because cilia distal segments are truncated in *osm-6* loss-of-function mutants (35).

Since vertebrate CNG channels have a central role in light detection and photoreceptor calcium homeostasis, the CNG distribution defect observed in *C. elegans cfh-1* mutant sensory neurons might provide insight into CFH function in human photoreceptors. Supporting this idea, photoreceptors from individuals homozygous for the AMD high-risk CFH Y402H variant showed increased relative levels of CNGA1 and CNGB1 subunits located in photoreceptor outer segments (Figs 4, 5). This may have significant implications for retina health, as mutations in CNG channels that are thought to affect intracellular calcium levels, guanylate cyclase activating protein (GCAP) activity, and cGMP production have been linked to retinitis pigmentosa and AMD (36-38).

**Figure 5.**
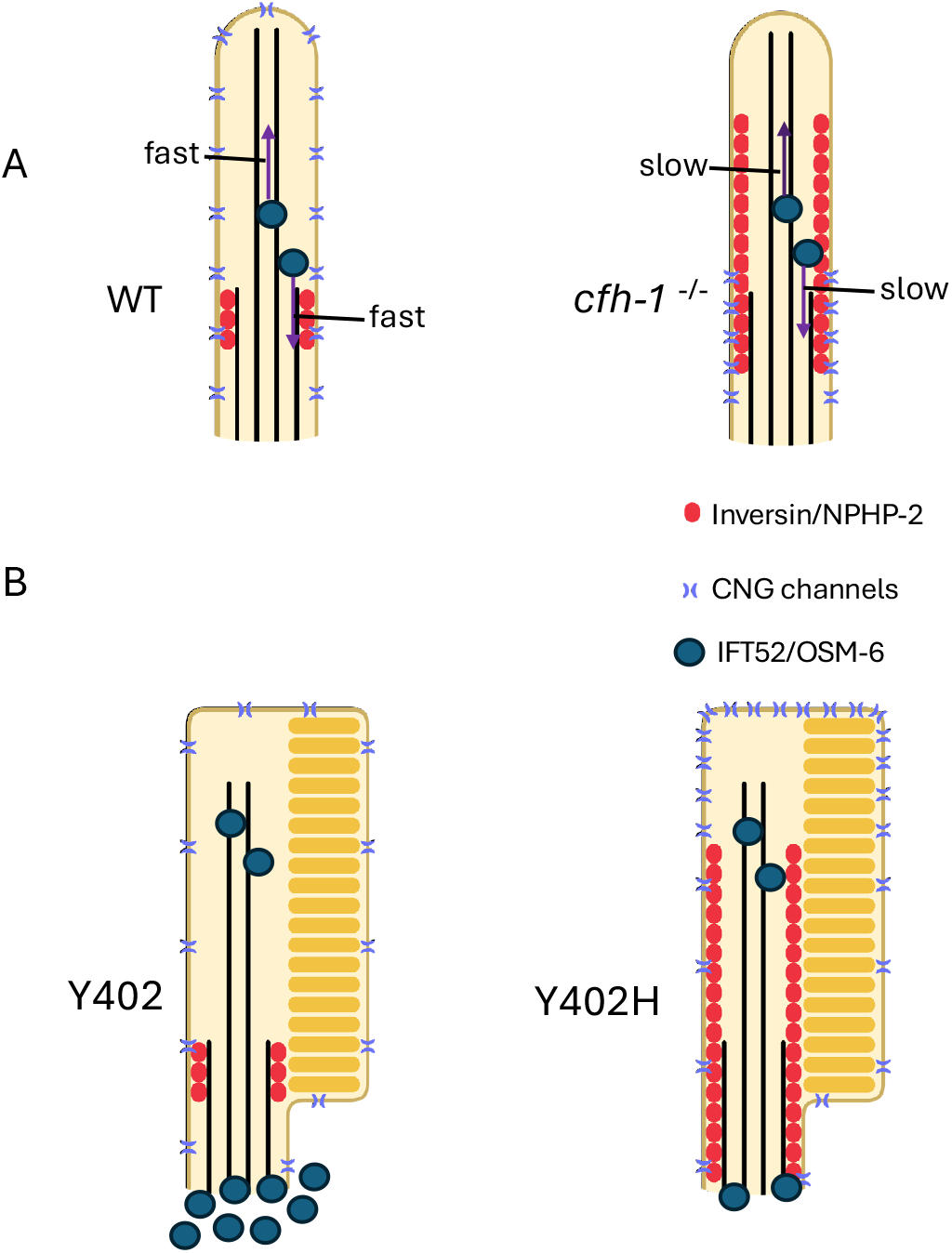
Schematic diagram summarizing relative locations. *Schematic diagram summarizing IFT52/OSM-6, CNG channel, and inversin/NPHP-2 distribution in distal compartments of WT and cfh-1 mutant C. elegans phasmid cilia (A) and CFH Y402 AMD low-risk Y402 and CFH Y402H AMD high-risk Y402H human photoreceptors (B)*. This figure summarizes data shown here and in our previous work (15). Note that inversin/NPHP-2 is diffuse throughout the cilia distal region in *C. elegans cfh-1* mutants and in AMD high risk photoreceptors. CNG channels are restricted to the middle segment in *cfh-1* mutant animals compared to WT animals but more concentrated in the photoreceptor outer segments in AMD Y402H high-risk photoreceptors compared to the photoreceptors of low-risk individuals. IFT52/OSM-6 anterograde and retrograde transport is slow in *cfh-1* mutant animals (purple arrows) and has decreased accumulation in the inner segments of AMD high-risk Y402H photoreceptors.

### Potential implications for photoreceptor dysfunction and retina diseases

Thinning of the photoreceptor layer is associated with AMD high-risk CFH variants and is one of the earliest AMD biomarkers, appearing decades before other clinical symptoms (10). Although it has been suggested that thinning may be a result of a defect in cholesterol recycling, there may be alternative models worth considering based on the abnormal IFT52/OSM-6 distribution demonstrated here. *C. elegans* IFT52/OSM-6 mutants have shortened axonemes presumably because of defective delivery of cargo needed for maintenance of axoneme length (35). Similarly, mutations in IFT52/OSM-6 result in shortening of cilia and retinal degeneration in humans (39). It therefore seems plausible that a defect in IFT52/OSM-6 transport in CFH loss-of-function photoreceptors could contribute to PS thinning due to a disruption of the equilibrium between the delivery of cilia structural proteins to photoreceptor outer segments and the absorption of shed material by retinal pigmented epithelial (RPE) cells (10).

In addition to photoreceptor thinning, another early biomarker linked to AMD high-risk variants is delayed rod-mediated dark adaptation (RMDA) (11). Although current models suggest that delayed RMDA may be due to a defect in retinoid transport to photoreceptors from choroid vasculature and RPE cells (40), the mechanistic connections between CFH and retinoid transport is not clear. Based on the preferential localization of CFH on rods, which are primarily responsible for dark adaptation, coupled with the CFH-dependent IFT52/OSM-6 and CNG distribution defects shown here, it may be reasonable to speculate that mis-localization of visual cycle components, including CNG, that affect intracellular calcium and cGMP levels, may contribute to delayed RMDA and other early manifestations of AMD.

Interestingly, a recent study demonstrated that regions of the retina that demonstrate delayed RMDA coincide with regions where the integrity of the interdigital zone (IZ), where photoreceptor outer segments overlap with apical processes of RPE cells, is compromised (41). The overlap in RMDA and IZ defects suggest that these phenomena share a common mechanism (41). It may be worth considering that a defect in IFT mediated transport of visual cycle and structural components within photoreceptors may contribute to this putative mechanism.

## Material and Methods

### C. elegans Strains and microscopy

A list of the *C. elegans* strains used in this work is included in Table S1.

Adult C. elegans strains were immobilized for 10 minutes with 15 µm levamisole (LKT laboratories) in M9 buffer and placed on a 5% agar pad on a microscope slide. Imaging was performed with a Nikon A1R confocal microscope with Nikon Elements software.

### Timelapse Live imaging, kymograph generation and analysis

#### Sample Preparation

To synchronize the ages of C. elegans, hermaphrodites at the L4 stage were transferred to fresh nematode growth medium (NGM) plates seeded with E. coli OP50. After two days incubation, adult worms were transferred to new plates on adult day 2 to separate them from younger worms, ensuring the collection of synchronized adult day 4 worms. On adult day 4, the worms were anesthetized and immobilized in 3 µl of 10 mM levamisole in M9 buffer on a 5% agar pad placed on a microscope slide, which was covered with a glass coverslip.

#### Imaging Setup

Phasmid cilia were imaged using a Nikon A1R confocal and super-resolution system equipped with a 100X objective lens with a Nipkow spinning disk. Time-lapse images were acquired for 30 seconds with an exposure time of 200 ms. A total of 150 images were collected and stacked for further analysis.

#### Kymograph Generation and Analysis

The stacked images were processed in ImageJ using the macro tool Kymograph Clear 2.0 (42) to generate kymographs. Kymograph Clear 2.0 automatically distinguishes moving particles based on their direction and generates kymographs for each set of directional tracks, as described previously (42). Particle tracks on the kymographs were manually traced. The generated kymographs were further analyzed with Kymograph Direct (42). The velocities of the traced particles in each direction were measured at positions along the cilium. Velocity values were compiled at 0.1 µm intervals from the ciliary base and averaged for downstream analysis.

### CFH custom antibody

Mice were immunized subcutaneously to 50 μg purified human Factor H (Sigma) emulsified in complete Freund’s Adjuvant and boosted two weeks later with 25 μg in incomplete Fruend’s Adjuvant. Two weeks later, mice were bled from the facial vein and sera were tested by ELISA on the immunogen. Microtiter wells were coated with 1 μg/ml factor H in phosphate-buffered saline (PBS) overnight and blocked for 2 hours with 1% casein. The results of the antisera were compared to pre-immune sera and tested in Western Blots (Fig. S3).

### Human stem cell maintenance and retinal organoid differentiation

Wildtype WA09 cells were maintained in mTeSR™1 (85857, StemCell Technologies) on 1% (v/v) Matrigel-GFR™ (354230, BD Biosciences) coated plate. Cells were maintained at 37ºC in a HERAcell 150i or 160i at 10% CO2 and 5% O2 (Thermo Fisher Scientific). Cells were then passaged and maintained following Wahlin et. al (43).

Human retinal organoids were generated with the AMASS (agarose microwell array seeding and scraping) protocol (44) with the minor alterations. Wells of an AggreWell™800 plate (34860, Stem Cell Technologies) were prepared by adding 500uL of rinsing solution to each well and spun at 2000xg for 5 minutes. Each well was rinsed with 2mL mTeSR™1. 1mL of mTeSR™1 plus 5 µM Blebbistatin (B0560, Sigma) was added to each well and put in hypoxic incubator to stay warm. Stem cell were washed with 2mL metal-free 1XPBS (14190094, Gibco), and incubated for 5 minutes at 37°C in 600uL 0.5mM RT EDTA (15575020, Invitrogen, diluted in metal-free 1XPBS). After removing the EDTA, 1mL of warm accutase (SCR005, Sigma) was added to the cells and incubated for 10 minutes at 37°C. 2mL of warmed mTeSR™1 was added to stop the reaction, and pipetted a few times to get to single cell solution. The suspension was spun for 5 minutes at 0.7 rpm, and the pellet resuspended in 500uL mTeSR™1 + blebbistatin for counting. Wildtype WA09 stem cells were seeded at 700 cells per microwell. Each well was filled to contain a total of 1.5ml cell suspension/mTeSR™1/blebbistatin and left in a hypoxic incubator overnight. The AMASS protocol was then followed (44) until organoids were 190 days old.

### Immunohistochemistry of human retinal organoids

190 day old retinal organoids were fixed at room temperature in fresh 4% formaldehyde and 5% sucrose in 1XPBS for 50 minutes. Organoids were rinsed 3X in 5% sucrose in 1XPBS, then incubated in a sucrose gradient for 30 min intervals at 4°C (6.75% followed by 12.5%). Finally organoids were incubated for 2 hours-overnight in 25% sucrose at 4°C. Organoids were incubated for 2 hours-overnight at 4°C in blocking solution (0.2% Triton X100, 4% donkey serum in 1XPBS). Organoids were incubated with the following primary antibodies in blocking solution for 16-36 hours at 4°C: custom mouse anti-CFH (1:50) chick anti-SWopsin (1:200) and chick anti-LMopsin (1:200, both gift from Jeremy Nathans) and rabbit anti-Rhodopsin (1:250, PA5-85608, Invitrogen). Organoids were washed 3X for 15 minutes in 1XPBS, and then incubated with the following secondary antibodies diluted 1:400 in blocking solution overnight at 4°C: Alexa Fluor-conjugated donkey anti-mouse 555, donkey anti-chick 488 and donkey anti-rabbit 647 (Molecular Probes). Organoids were washed 3X for 10 minutes in 1XPBS. Organoids were counterstained with Hoechst 33342 nuclear marker (1:2000 in 1XPBS) for 10 min and washed 3X for 15 min in 1XPBS. Organoids were mounted for imaging in Slow Fade Gold (S36940, Thermo Fisher Scientific).

### Imaging and analysis of CFH in human retinal organoids

Three 0.25mm^2^ random regions of interest were imaged from 3 different organoids with a laser scanning confocal Zeiss LSM 980 inverted microscope using a Plan-Apochromat 63x/1.4 Oil DIC M27 objective at 2% laser power for each channel. The opsin staining labels both the outer segments and somas of the photoreceptors. Mature rod outer segments are extremely long and our organoids are rod-enriched, with neighboring rod outer segments often overlaying each other. This caused technical difficulties in confidently determining the proportion of photoreceptors expressing CFH. To avoid this issue, we imaged and assessed CFH expression within rod and cone somas only. During imaging, we manually centered 4 µm Z-stacks with 1µm intervals over photoreceptor somas.

CFH expression in rod and cone somas was analyzed using the HALO Image Analysis Platform 3.5.3577 (Indica Labs, Inc.). To calculate the proportion of photoreceptors expressing CFH, we used the Object Colocalization FL v2.1.4 module. Rod somas (Rhodopsin positive), cone somas (SWopsin or LW/MWopsin positive) and CFH protein (CFH positive) was identified. We counted the total number of rod and cone somas detected, and then determined how many of these overlap with CFH expression by > 10% of their total area. Average ratios of CFH-expressing photoreceptors were compared using a one-way ANOVA with Tukey’s multiple comparisons test (R Studio). GraphPad Prism version 9.3.1 (San Diego, California USA, www.graphpad.com) was used to generate graphs.

### Immunohistochemistry of mouse and human retina sections

Postmortem mouse and human eyes were embedded in paraffin and sectioned as described (30). Sectioned eyes were deparaffinized in Xylene and hydrated by washing in 100%, 95%, 70%, 50%, 30% Ethanol and H2O (5 mins each). After washing in Tris-buffered Saline with 0.1% tween for 5 mins and PBS 3 times for 5 mins, sections were blocked in 10% Donkey serum in PBS with 0.5% Triton-X-100 for 1 hour. Sections were incubated at 4°C overnight with the following primary antibodies in blocking solution: mouse monoclonal antibody to Complement factor H (1:50, custom antibody described above or Invitrogen C18/3), mouse monoclonal antibody to rhodopsin (4D2) (1:400, NBP2-59690, Novus Biologicals) polyclonal antibody to rhodopsin (1:200, PA5-85608, Invitrogen), rabbit polyclonal antibody to CNGA1 (1:200 PA5101516, Invitrogen), rabbit polyclonal antibody to CNGB1 (1:200, 30557-1-AP, Proteintech), rabbit polyclonal antibody to IFT52 (1:10, 17534-1-AP, Proteintech), Goat polyclonal antibody to IFT88 (1:200, PA5-18467, Invitrogen). After washing in PBS 3 times for 10mins each, sections were incubated with the following secondary antibodies for 1 hour in RT: Donkey anti-mouse Alexa Fluor 647 conjugated secondary antibody (1:200, A31571, Thermo Fisher Scientific),Donkey anti-rabbit Alexa Fluor 488 conjugated secondary antibody (1:200, A21206, Thermo Fisher Scientific), and Donkey anti-goat Alexa Fluor™ 555 secondary antibody (1:200, A21432, Thermo Fisher Scientific). After a rinse and 3 washes in PBS for 10 mins each, sections were counterstained by 5µM DAPI for 7min and washed in PBS 3 times for 20mins. Sections were mounted in Slow Fade Gold (S36936, Invitrogen by Thermo Fisher Scientific) or KPL mounting Medium (5570-0005, Seracare) and sealed by cover grass and nail polish. Slides were imaged using a Nikon A1R confocal microscope with Nikon Elements software.

## Statistical analysis

One-way ANOVA followed by Tukey’s or Dunnett’s multiple comparisons test and correlation analysis were performed using GraphPad Prism version 10.2.3 for macOS (GraphPad Software, San Diego, California USA, www.graphpad.com).

## Acknowledgments

The authors thank Maureen Barr, Inna Nechipurenko, and Piali Sengupta for their generosity in sharing *C. elegans* strains and constructs. We also thank Kevin Rossomando and Qiang Lin for technical assistance and Brad Shibata and the NEI Core Facilities grant P30-EY012576 (UCD) at UC Davis for mouse retina sections. J.A.M is supported by the Canadian Institutes of Health Research. Some *C. elegans* strains were provided by the CGC, which is funded by NIH Office of Research Infrastructure Programs (P40 OD010440) and the C. elegans Reverse Genetics

Core Facility at the University of British Columbia, which is part of the international C. elegans Gene Knockout Consortium. Research reported in this publication was supported by the National Eye Institute of the National Institutes of Health under Award Number R01EY032868 to B.E.V. The content is solely the responsibility of the authors and does not necessarily represent the official views of the National Institutes of Health.

## Figure Legends

Figure S1. *Comparison of anterograde IFT of dynein light-intermediate chain XBX-1 and IFT-A component IFT140/CHE-11 in WT and cfh-1 mutant C. elegans day 4 adults.*

A, Plots of anterograde IFT velocity vs. distance from cilia base for dynein light-intermediate chain XBX-1 (left) and IFT-A component IFT140/CHE-11 in phasmid sensory neurons of WT(black) and *cfh-1* mutant (yellow) animals. Note that IFT velocities for IFT140/CHE-11 are similar in WT and *cfh-1(em14)* animals while XBX-1 IFT is faster in *cfh-1* mutant animals than in WT. B, Representative kymograph plots upon which the data are based. Horizonal scale bars represent 1µm and vertical scale bars indicate a 5 second interval.

Figure S2. NPHP-2 does not affect IFT rates.

A, *Comparison of anterograde and retrograde IFT of IFT52/OSM-6 in WT and nphp-2(gk653) mutant C. elegans day 1 adults*. Note that there is little or no difference in IFT52/OSM-6::GFP IFT in WT and *nphp-2* mutant animals. B, *Comparison of anterograde and retrograde IFT of IFT52/OSM-6 in WT, cfh-1(em14) and nphp-2 (gk653); cfh-1(em14) double mutant day 4 adults*. Note that there is little or no difference in IFT52/OSM-6::GFP IFT in *cfh-1* and *cfh-1;nphp-2* mutant animals.

Figure S3. CFH expression associated with human rod and cone photoreceptors

A, Immunofluorescence with antibodies to CFH protein (magenta) and rhodopsin (cyan) in left panel or SWopsin and LWMWopsin (green, right panel) in 190 day old human retinal organoids. White arrows point to examples of rods with CFH. Right, scatter plot showing that a significantly higher proportion of rods contain CFH compared to cones in human organoids. Asterisks indicate significant differences detected by an ANOVA followed by a Tukey HSD test, where *** = p < 0.001. Error bars represent standard deviation of the mean. B, Antibody staining of human retina sections reveals CFH staining of photoreceptor outer segments. Also show is rhodopsin staining to indicate rod outer segments (brackets). Scale bar is 5µm. C, Western Blot of human plasma using commercial and custom mouse antibodies to human CFH

