## Supplemental Table 1 for "Complement Factor H and its *C. elegans* homolog regulate IFT52/OSM-6 and CNG channel localization in sensory neurons"

Supplemental Table S1 List of strains used

| STRAIN | GENOTYPE |
| --- | --- |
| BT61 | *myEx1[Posm-5::OSM-5::GFP + pRF4];* [*xbx-1*](https://cgc.umn.edu/strain/search?st1=xbx-1&sf1=all)*(*[*cas502*](https://cgc.umn.edu/strain/search?st1=cas502&sf1=all)*[*[*xbx-1*](https://cgc.umn.edu/strain/search?st1=xbx-1&sf1=all)*::[tagRFP](https://cgc.umn.edu/strain/search?st1=tagRFP&sf1=all)])* |
| BT63 | *cfh-1 (em14); myEx1[Posm-5::OSM-5::GFP + pRF4];* [*xbx-1*](https://cgc.umn.edu/strain/search?st1=xbx-1&sf1=all)*(*[*cas502*](https://cgc.umn.edu/strain/search?st1=cas502&sf1=all)*[*[*xbx-1*](https://cgc.umn.edu/strain/search?st1=xbx-1&sf1=all)*::[tagRFP](https://cgc.umn.edu/strain/search?st1=tagRFP&sf1=all)])* |
| BT64 | *mnIs17 [osm-6::GFP + unc-36(+)];* [*xbx-1*](https://cgc.umn.edu/strain/search?st1=xbx-1&sf1=all)*(*[*cas502*](https://cgc.umn.edu/strain/search?st1=cas502&sf1=all)*[*[*xbx-1*](https://cgc.umn.edu/strain/search?st1=xbx-1&sf1=all)*::[tagRFP](https://cgc.umn.edu/strain/search?st1=tagRFP&sf1=all)])* |
| BT65 | *cfh-1 (em14); mnIs17 [osm-6::GFP + unc-36(+)];* [*xbx-1*](https://cgc.umn.edu/strain/search?st1=xbx-1&sf1=all)*(*[*cas502*](https://cgc.umn.edu/strain/search?st1=cas502&sf1=all)*[*[*xbx-1*](https://cgc.umn.edu/strain/search?st1=xbx-1&sf1=all)*::[tagRFP](https://cgc.umn.edu/strain/search?st1=tagRFP&sf1=all)])* |
| BT66 | *myEx[CHE-11::GFP+pRF4];* [*xbx-1*](https://cgc.umn.edu/strain/search?st1=xbx-1&sf1=all)*(*[*cas502*](https://cgc.umn.edu/strain/search?st1=cas502&sf1=all)*[*[*xbx-1*](https://cgc.umn.edu/strain/search?st1=xbx-1&sf1=all)*::[tagRFP](https://cgc.umn.edu/strain/search?st1=tagRFP&sf1=all)])* |
| BT67 | *cfh-1 (em14); myEx[CHE-11::GFP+pRF4];* [*xbx-1*](https://cgc.umn.edu/strain/search?st1=xbx-1&sf1=all)*(*[*cas502*](https://cgc.umn.edu/strain/search?st1=cas502&sf1=all)*[*[*xbx-1*](https://cgc.umn.edu/strain/search?st1=xbx-1&sf1=all)*::[tagRFP](https://cgc.umn.edu/strain/search?st1=tagRFP&sf1=all)])* |
| BT68 | *byEX836[odr-4p::tax-4::GFP + myo-2p::mCherry];* *emEX[bbs-8p::NPHP-2::mCherry+pRF4]* |
| BT69 | *cfh-1(em14); byEX836[odr-4p::tax-4::GFP + myo-2p::mCherry];* *emEX[bbs-8p::NPHP-2::mCherry+pRF4]* |
| BT70 | *nphp-2(gk653*); *emIs20[bbs-8p::OSM-6::GFP+pRF4]* |
| BT71 | *cfh-1(em14); nphp-2(gk653);* *emIs20[bbs-8p::OSM-6::GFP+pRF4]* |
