## Supplementary figures and images for "Complement Factor H and its *C. elegans* homolog regulate IFT52/OSM-6 and CNG channel localization in sensory neurons"

### Supplemental Figures

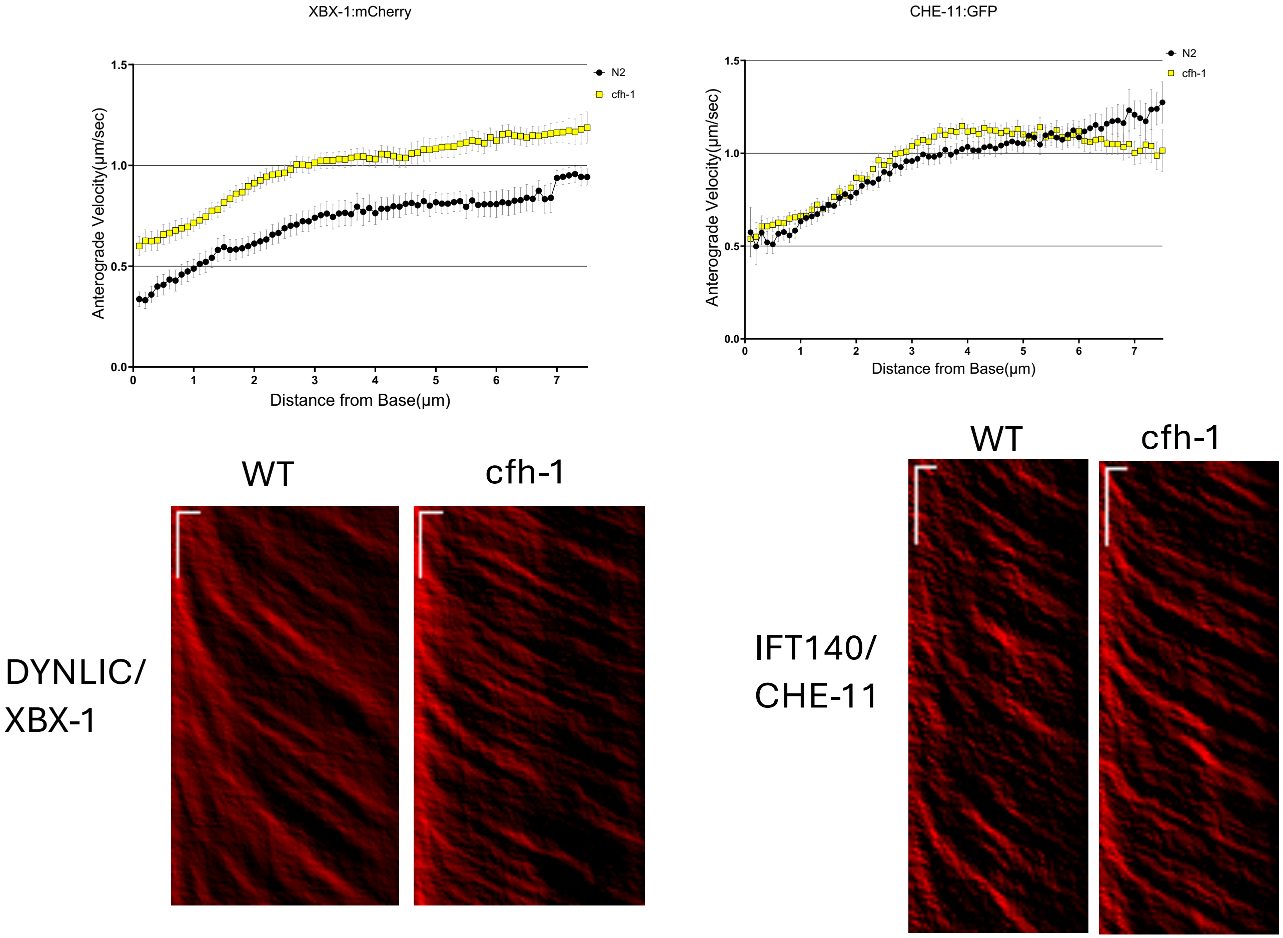

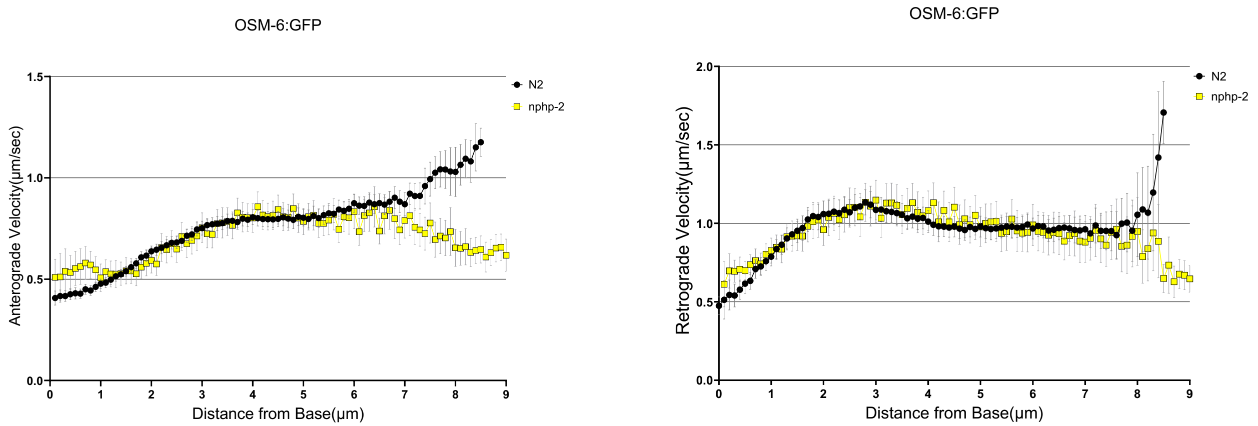

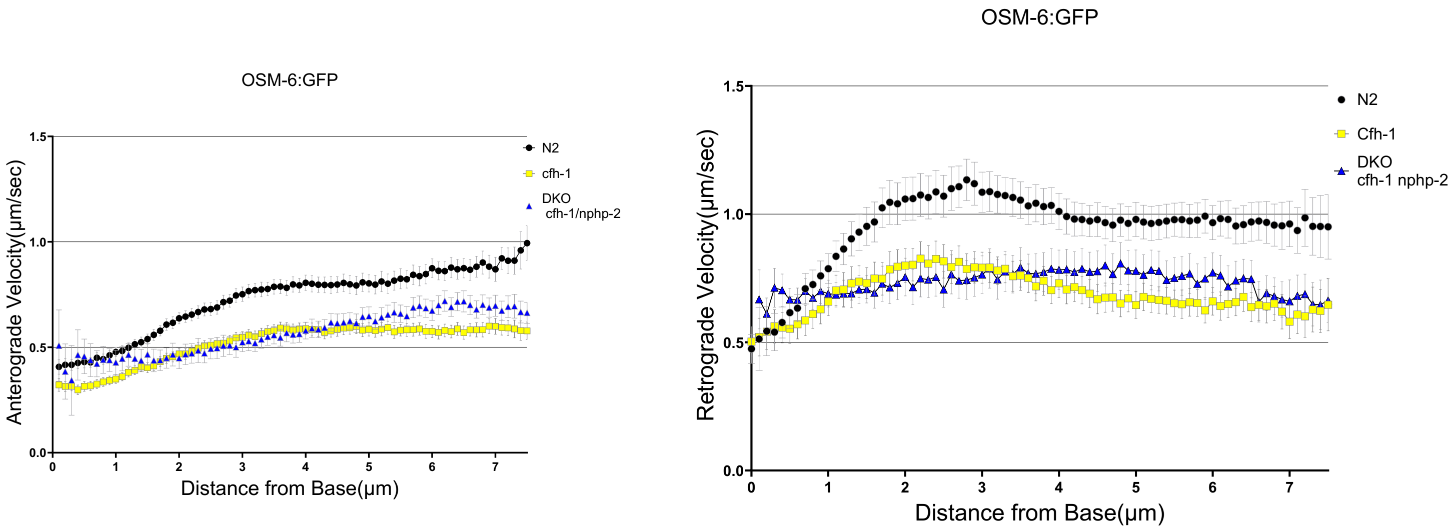

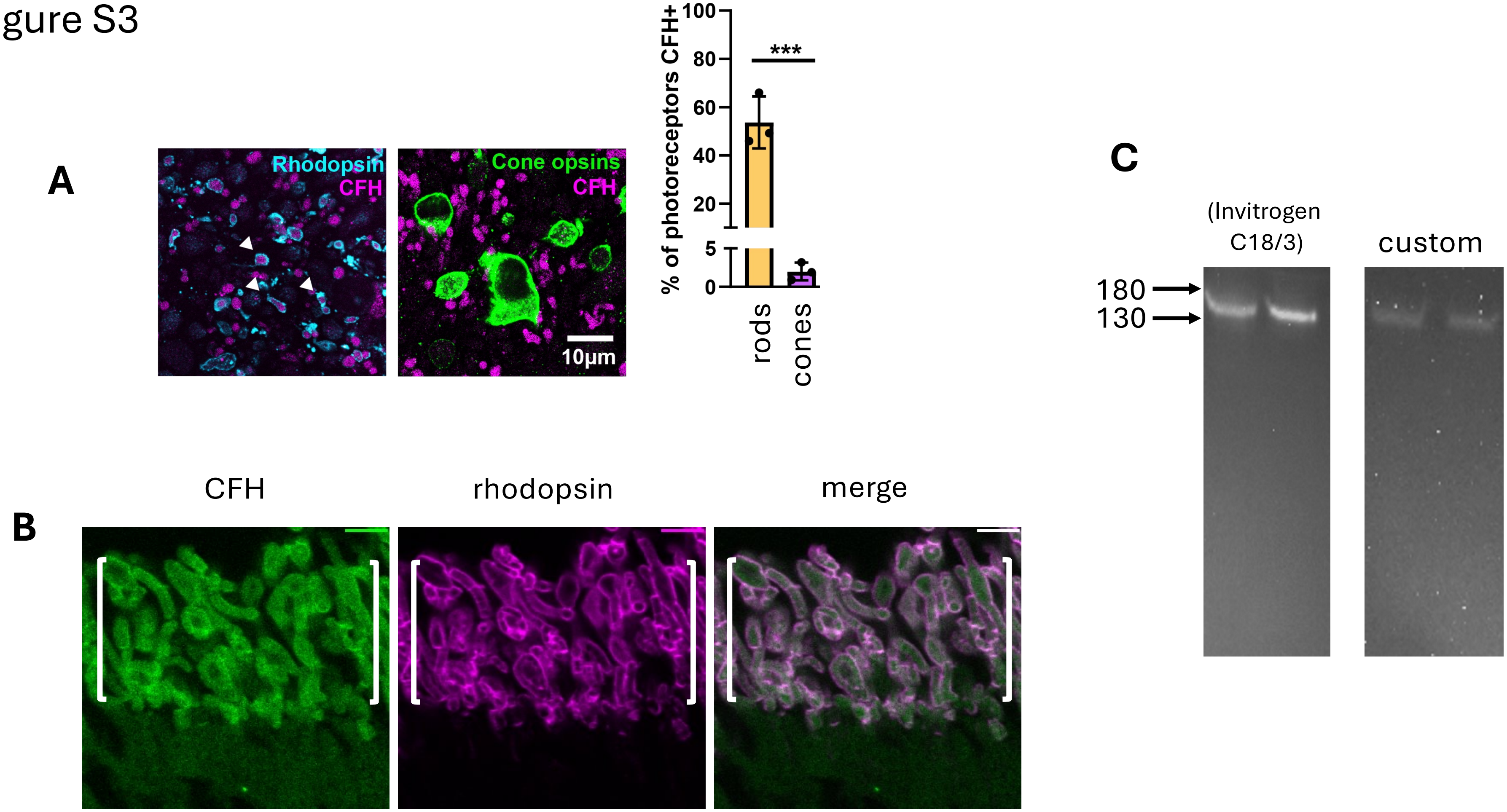
